# 3D Chromatin Structure in Chondrocytes Identifies Putative Osteoarthritis Risk Genes

**DOI:** 10.1101/2022.05.16.492146

**Authors:** Eliza Thulson, Eric S. Davis, Susan D’Costa, Philip R. Coryell, Nicole E. Kramer, Karen L. Mohlke, Richard F. Loeser, Brian O. Diekman, Douglas H. Phanstiel

## Abstract

Genome-wide association studies (GWAS) have identified over 100 loci associated with osteoarthrtis (OA) risk, but the majority of OA risk variants are non-coding, making it difficult to identify the impacted genes for further study and therapeutic development. To address this need, we used a multi-omic approach and genome editing to identify and functionally characterize potential OA risk genes. Computational analysis of GWAS and ChIP-seq data revealed that chondrocyte regulatory loci are enriched for OA risk variants. We constructed a chondrocyte specific regulatory network by mapping 3D chromatin structure and active enhancers in human chondrocytes. We then intersected these data with our previously collected RNA-seq dataset of chondrocytes responding to fibronectin fragment (FN-f), a known OA trigger. Integration of the three genomic datasets with recently reported OA GWAS variants revealed a refined set of putative causal OA variants and their potential target genes. One of the novel putative target genes identified was *SOCS2*, which was connected to a putative causal variant by a 170 Kb loop and is differentially regulated in response to FN-f. CRISPR-Cas9-mediated deletion of *SOCS2* in primary human chondrocytes from three independent donors led to heightened expression of inflammatory markers after FN-f treatment. These data suggest that *SOCS2* plays a role in resolving inflammation in response to cartilage matrix damage and provides a possible mechanistic explanation for its influence on OA risk. In total, we identified 56 unique putative OA risk genes for further research and potential therapeutic development.

## INTRODUCTION

Osteoarthritis (OA) affects over 300 million people world-wide, yet treatment options are limited in large part because the mechanisms driving OA are not fully understood ^1,2^. Genome-wide association studies (GWAS) have identified over 100 loci associated with OA risk^3^, but translating these broad loci into therapeutic targets has been challenging for several reasons. First, the effects of disease-associated variants are likely cell-type and context specific^4^; therefore, studying these variants in the correct system that mimics the OA phenotype is required. Second, linkage disequilibrium (LD) between nearby variants makes it difficult to identify the causal variant(s) at each locus. Finally, because the majority of OA risk variants occupy non-coding regions of the human genome and can regulate genes up to a million base pairs away, the genes impacted by most OA risk variants are unknown.

Several studies have successfully used genomic and bioinformatic techniques to identify the genes impacted by gene-distant non-coding GWAS variants for a variety of disease phenotypes^5–9^. Mapping regulatory loci using ChIP-seq, ATAC-seq, or CUT&RUN and intersecting the resulting data with disease-associated variants can identify a short list of putative causal variants. These variants can then be linked to potential target genes by quantifying 3D chromatin contacts using Hi-C or other chromatin conformation capture (3C) techniques. For example, chromatin interaction data was used to determine that an obesity-associated variant located in an intron of the *FTO* gene affects expression of the downstream genes *IRX3* and *IRX5*, which are involved in obesity-related biological processes^5^. Likewise, Hi-C in human cerebral cortex identified FOXG1 as a distal target of a schizophrenia GWAS variant, supporting its potential role as a schizophrenia risk gene^6^.

Because the effects of disease-associated variants are likely limited to particular biological states^4^, studies of their impact must be conducted in the correct cellular and biological context. Several pieces of evidence suggest that chondrocytes—particularly those responding to cartilage matrix damage—are one of the most likely cell types to be affected by OA risk variants. Cartilage breakdown and loss is a primary feature of OA. Chondrocytes are the only cell type found in cartilage and are responsible for maintaining the cartilage matrix. Osteoarthritic cartilage harbors activated chondrocytes that exhibit a proinflammatory phenotype thought to contribute to progressive cartilage degradation, which includes production of bioactive matrix fragments^10,11^. We have developed an *ex vivo* system that simulates the OA chondrocyte phenotype by treating primary human articular chondrocytes with fibronectin fragment (FN-f)^12–15^. Fibronectin is a ubiquitous extracellular matrix protein, and high levels of FN-f are present in cartilage and synovial fluid of OA joints^16,17^. Subsequently, FN-f has been shown to be an OA mediator that recapitulates gene expression changes associated with OA^12–14,18^. We have leveraged this model of OA for use in clonal populations of genome-edited primary human chondrocytes, allowing us to quantify the phenotypic impact of putative target genes of genomic variants in an appropriate disease context.

In this study, we generated CUT&RUN and Hi-C datasets in human chondrocytes and intersected them with publicly available data from our lab and others. In doing so, we identified 56 putative OA risk genes, including *SOCS2*, whose promoter loops to an OA GWAS variant ~174 Kb away. Deletion of *SOCS2* in primary human chondrocytes using CRISPR-Cas9 led to heightened expression of inflammatory markers in response to treatment with FN-f, providing a possible mechanism for influencing OA risk.

## RESULTS

### OA risk variants are enriched in chondrocyte regulatory loci

One of the first steps in decoding GWAS variant mechanisms is to determine the cell types that are likely mediating genetic OA risk. While different risk variants may impact distinct cell types, one approach to help direct research is to determine the cell types which harbor regulatory loci (e.g. enhancers) that are enriched for risk variants. To accomplish this, we performed SNP enrichment analysis using the Genomic Regulatory Elements and Gwas Overlap algoRithm (GREGOR)^19^. Publicly available H3K27ac, H3K4me1, and H3K4me3 ChIP-seq peaks from the NIH Roadmap Epigenomics Mapping Consortium (Roadmap) were merged to define regulatory elements for 98 cell types. GREGOR was used to determine each cell type’s enrichment for 104 OA GWAS signals recently published in Boer et al.^2^

The regulatory elements of “Chondrocytes from Bone Marrow Derived Mesenchymal Stem Cell Cultured Cells” (E049) exhibited a strong effect size and p-value of enrichment for OA risk variants (**Fig. 1A**), suggesting that many OA risk variants may impact regulatory events in chondrocytes. This is consistent with the known role of chondrocytes in maintaining joint homeostasis. Chondrocytes have been heavily implicated in OA, as activation of chondrocytes by mechanical and inflammatory stimuli triggers downstream inflammatory and catabolic response pathways in diseased tissue^20–23^. An example of an OA risk variant that overlaps a chondrocyte-specific regulatory element (H3K27ac peak) is shown in **Figure 1B**. For comparison, **Figure 1C** shows a non-OA associated variant that overlaps a non-cell type specific, or ubiquitous, enhancer on chromosome 10 that is active in >90% of the 98 cell types evaluated. These examples underscore the importance of interpreting GWAS risk variants in light of the correct cellular context, as the variant-H3K27ac peak overlap shown in **Figure 1B** would not have been detected in any of the other cell types investigated. In addition to E049, IMR90 fetal fibroblasts (E017) and HSMM cell derived Skeletal Muscle Myotubes (E121) were also enriched, suggesting that OA risk variants may also contribute to disease risk through altering the function of fibroblasts and muscle. However, given the strong enrichment in chondrocytes and their documented role in OA biology, we chose to focus our investigation of OA GWAS variants in human chondrocytes.

**Figure 1.**
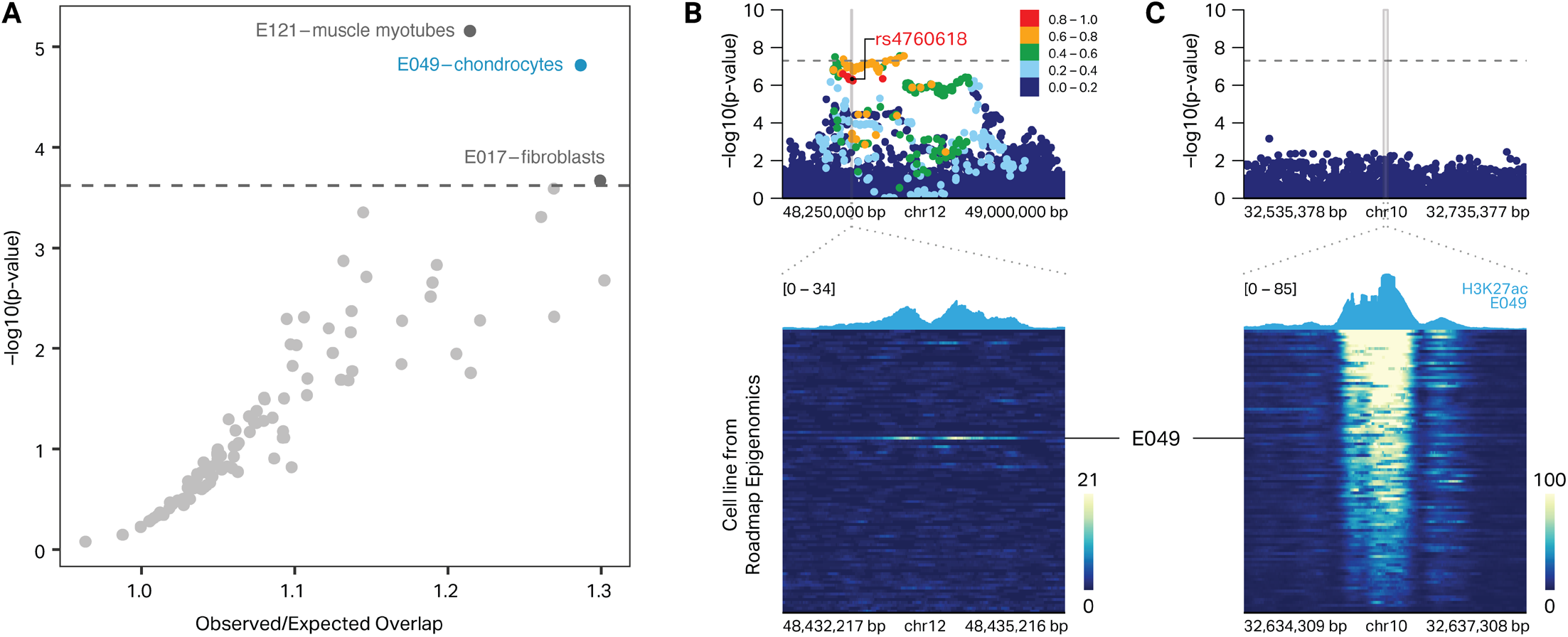
OA risk variants are enriched in chondrocyte regulatory elements. **(A)** Enrichment analysis of 98 cell types from the NIH Roadmap Epigenomics Mapping Consortium reveals that OA GWAS variants are enriched in the regulatory regions (H3K27ac, H3K4me1, or H3K4me3 ChIP-seq peaks) of chondrocytes, skeletal muscle myotubes, and fibroblasts. **(B)** Heatmap of H3K27ac signal from 98 cell types (bottom) high-lights a chondrocyte-specific enhancer that overlaps Knee/Hip osteoarthritis risk variant2 (rs4760618, circled) that is in high LD (r2 > 0.8) with the lead variant (rs7967762, red diamond) for this locus (top). **(C)** Heatmap of H3K27ac signal from 98 cell types (bottom) highlights a ubiquitous enhancer (active in >90% of cell types) that does not overlap an OA GWAS variant (top).

### Multi-omic integration identifies putative variant-gene associations in OA

Due to high LD between variants and the fact the most risk variants reside in non-coding sequences, determining the causal variants and genes they impact remains a major challenge. To address these issues, we generated novel maps of epigenetic features in human chondrocytes and integrated them with GWAS results and publicly available genomic datasets to identify putative variant-gene associations for OA.

First, we identified OA risk variants that are predicted to directly affect protein sequences. We used ENSEMBL’s Variant Effect Predictor (VEP) tool to predict the consequences of 1,259 putative OA risk variants that were in high LD (r^2^> 0.8) with 104 OA GWAS signals from Boer et al^2^. VEP identified 29 variants at 19 loci predicted to affect the coding sequence of 24 unique genes (**Fig. 2A, top**). 18 of these variants encode a missense mutation impacting 17 genes, while 11 variants encode a synonymous mutation impacting 8 genes (**Fig. 2A, top**). Though synonymous variants do not impact the protein sequence directly, differences in transcription efficiency, tRNA availability, and mRNA stability introduced through these variants could contribute to the OA phenotype^24,25^. Of the 24 genes identified here, 6 exhibited differential expression in response to FN-f, (**Fig. 2A bottom**). Several of the genes identified have been previously implicated in OA, including Interleukin 11 (*IL11*), Solute Carrier Family 39 Member 8 (*SLC39A8/ZIP8*), and Serpin Family A Member 1 (*SERPINA1*). IL11 plays a role in bone turnover and is upregulated in subchondral bone and articular cartilage from OA tissue^26^. SLC39A8 is upregulated in OA chondrocytes and suppression of SLC39A8 in a mouse OA model significantly reduces cartilage degradation^27^. SERPINA1, a serine protease inhibitor with anti-inflammatory capabilities^28^, is downregulated in OA^29,30^.

**Figure 2.**
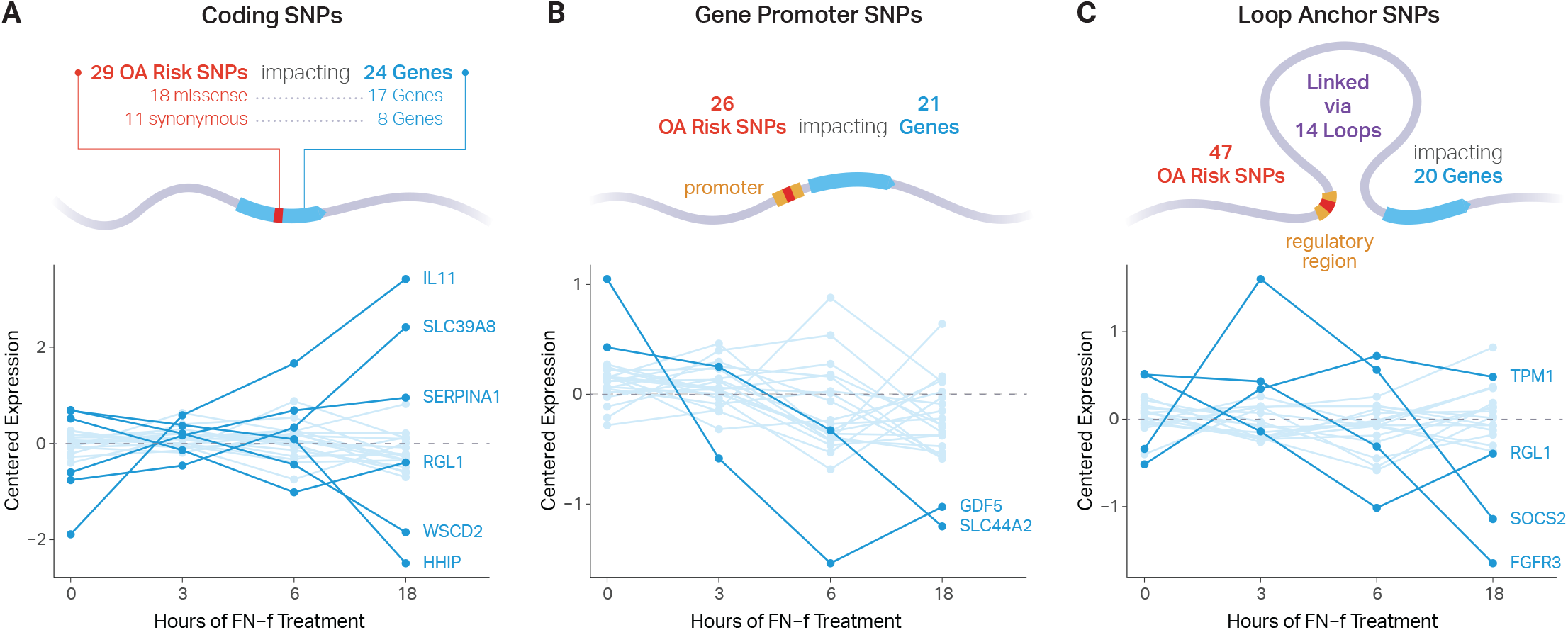
Multi-omic integration for assigning SNPs to putative OA risk genes. **(A)** ENSEMBL’s Variant Effect Predictor tool identified 29 unique OA risk SNPs (18 missense and 11 synonymous) overlapping coding regions of 24 unique genes (17 missense, 8 synonymous). **(B)** 26 unique OA risk SNPs overlapped both a chondrocyte regulatory region (H3K27ac, H3K4me1, or H3K4me3 ChIP-seq peaks) and a gene promoter for 21 unique genes. **(C)** 47 unique SNPs overlapped chondrocyte regulatory regions connected to 20 unique gene promoters via 14 C-28/I2 chromatin loops. RNA-seq data from our ex-vivo OA model depicts how putative OA risk genes change in response to FN-f. Normalized expression of genes are shown below each category over an 18 hour time-course of fibronectin fragment (FN-f) treatment. Differential genes (p ≤ 0.01, absolute log2 fold-change ≥ 1.25) are colored and labeled.

Next, we identified OA risk variants that could impart their phenotypic impact by altering promoter and/ or enhancer activity. We used CUT&RUN to map histone H3K27 acetylation (H3K27ac)—a mark of active enhancers and promoters—in primary human chondrocytes isolated from the knees of 2 cadaveric donors. We called H3K27ac peaks and merged them with all available marks (including active and repressive marks) from the E049 chondrocyte cell line from Roadmap Epigenomics to define a set of chondrocyte regulatory elements. Out of 1,259 putative OA risk variants, this overlap identified 507 plausible regulatory variants.

Intersecting these 507 plausible regulatory variants with gene annotations (UCSC) identified 26 unique variants that overlapped the promoters of 21 genes (**Fig. 2B**). Two of these genes were differentially expressed in response to FN-f, both of which have been previously implicated in OA. Growth and Differentiation Factor 5 (GDF5), a member of the TGF-beta family, has roles in skeletal and joint development^31^ and has been identified as a major risk locus for OA^32,33^. Specifically, variants in the GDF5 enhancers *R4* and *GROW1* have been associated with altered anatomical features of the knee and hip, which are thought to confer an increased risk of OA^34–36^. Solute Carrier Family 44 Member 2 (SLC44A2, aka choline transporter-like protein 2) is a mitochondrial choline transporter that has been identified as an expression quantitative trait locus (eQTL) in OA tissue^37^ that colocalizes with the OA GWAS signal rs1560707^38^.

In addition to direct regulation of genes by their promoters, long-range regulation of genes also occurs via enhancer-promoter interactions mediated by chromatin loops^39^. To identify such connections, we conducted deeply sequenced (~2.8 billion reads) in situ Hi-C in C-28/ I2 chondrocyte cells and identified 9,271 chromatin loops with Significant Interaction Peak (SIP) caller^40^. To our knowledge, this is the first Hi-C map in a chondrocyte cell line, enabling us to discover novel OA-associated variant-gene connections. Overlapping these data with OA risk variants identified 14 unique loops that connected 47 unique variants among 14 loci to 20 unique gene promoters (**Fig. 2C**). Four of these genes were differentially expressed in response to FN-f (p ≤ 0.01, absolute log2 fold-change ≥ 1.25). Several of these genes have interesting implications for OA, including *FGFR3* (Fibroblast Growth Factor Receptor 3), which plays a role in skeletal development. FGFR3 may have an important function in the maintenance of articular cartilage^41–43^, possibly through the Indian hedgehog signaling pathway, which plays a role in regulating chondrocyte hypertrophy and the expression of cartilage matrix-degrading enzymes^44^. FGFR3 is also downregulated in OA tissues, further implicating its potential role in limiting articular cartilage degeneration^45,46^. All putative variant-gene associations are reported in **Supplementary Table 1**.

### Chondrocyte chromatin features identify *SOCS2* as a putative regulator of OA

Our multi-omic analysis identified an association between rs7953280 and the promoter of Suppressor Of Cytokine Signaling 2 (*SOCS2)*. rs7953280 is located in an intron of the *CRADD* gene, which is expressed at low levels in primary chondrocytes, does not change expression in response to FN-f, and lacks an obvious biological relevance to OA. However, rs7953280 overlaps a putative chondrocyte enhancer (i.e. histone H3K27ac peak), suggesting that it could alter the regulatory capacity of the enhancer and impact the expression of a proximal or distal gene. This enhancer is connected to the promoter of *SOCS2* via a 174 Kb chromatin loop (**Fig. 3A**). Unlike *CRADD, SOCS2’s* expression changes in response to FN-f, peaking at 3 hours (**Fig. 2C** and **3A orange signal tracks**). Moreover, *SOCS2* is known to play a role in resolving inflammatory response through NFKB and is downregulated in knee OA tissues^47,48^, making it an intriguing candidate as an OA risk gene. No other SNPs from this locus can be assigned to genes using our integrated approach.

**Figure 3.**
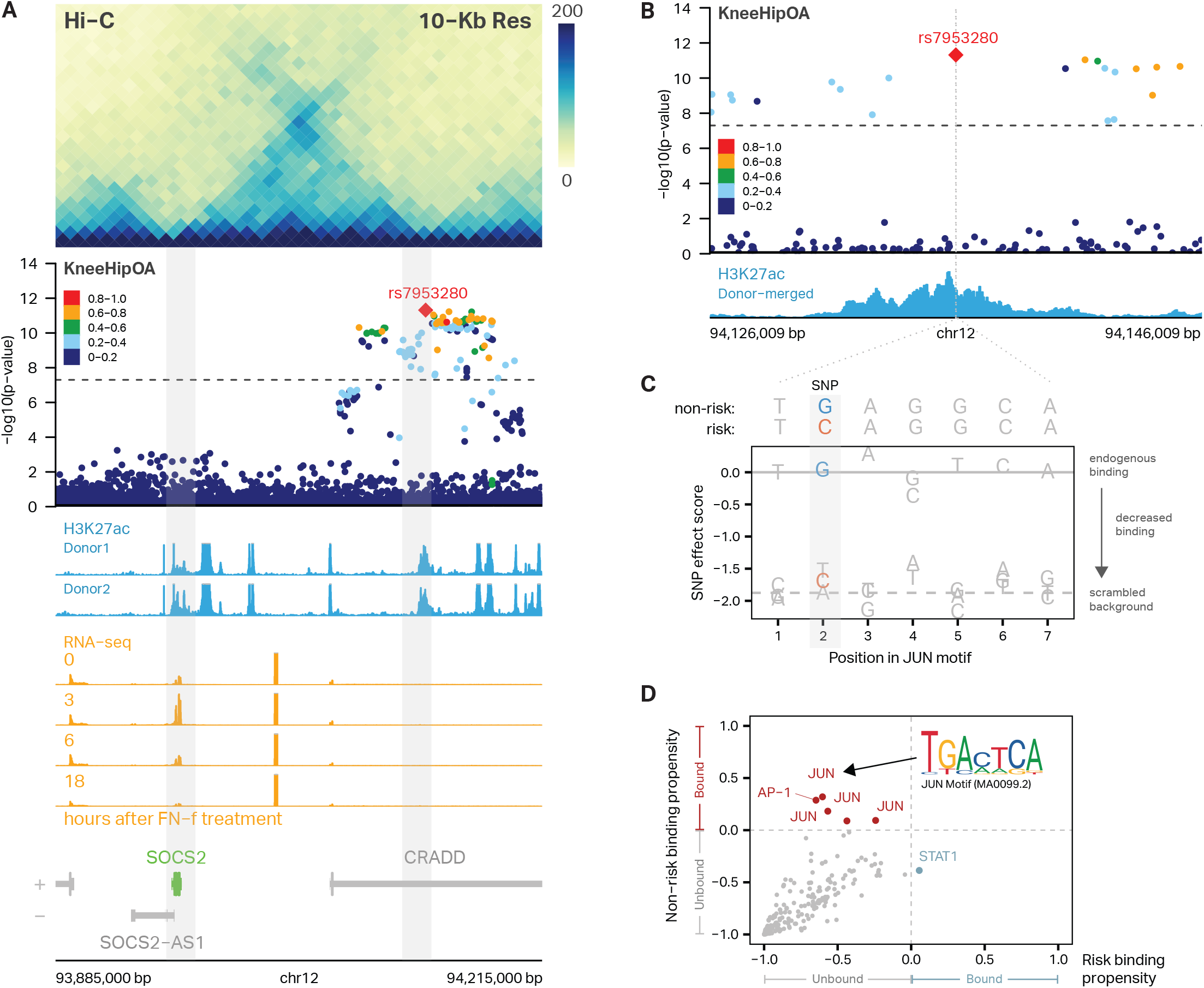
3D chromatin interactions identify SOCS2 as a putative regulator of OA. **(A)** Hi-C performed in C-28/I2 cells reveals a chromatin loop connecting OA risk variant rs7953280 (right gray bar) to the promoter of SOCS2 (left gray bar). rs7953280 is located in an intronic region of CRADD and overlaps an H3K27ac peak in primary articular human chondrocytes from two donors (blue signal tracks). SOCS2 is differentially expressed in response to treatment with FN-f. Gene tracks are shown below with +/-indicating gene strand. (B) Zoom-in on rs7953280 shows that the SNP is located within an H3K27ac peak in primary articular human chondrocytes. (C) Motif analysis identifies a JUN binding site at rs7953280. SNP effect matrix (SEM) data predicts decreased binding at the JUN motif (JASPAR ID: MA0099.2) with a G to C polymorphism in the second position. (D) Motif analysis from 211 precomputed SEMs from SEMpl predicts that JUN/AP-1 motifs (red, upper left quadrant) bind to the non-risk but not the risk allele.

To further understand how rs7953280 may confer risk for OA, we examined the sequence surrounding rs7953280 to see if it overlaps and alters any transcription factor (TF) binding motifs. Motif comparison with Tomtom from the MEME suite identified FOS and JUN as matching target motifs (**Fig. 3C, Supplementary Table 2**). FOS and JUN are members of the Activator Protein 1 (AP-1) complex, which is upregulated in response to FN-f^12^, and the inhibition of which prevents cartilage degradation in a model of OA^49–51^. We then used SNP effect matrices (SEMs) generated by the SNP effect matrix pipeline (SEMpl)^52^ to assess the predicted consequence of the G (non-risk) to C (risk) variant on binding of JUN or any other of the 211 motifs included with SEMpl (**Fig. 3D**). Most TFs are predicted to be unbound at both alleles. However, multiple JUN/AP-1 motifs are predicted to bind to the non-risk, but not the OA-risk sequence (**Fig. 3D**) providing further evidence that the G->C mutation in rs7953280 may disrupt JUN/AP-1 binding. Our analysis also showed that STAT-1 was predicted to bind only to the OA-risk sequence, although the SEM score was very close to the cutoff for predicted binding. Nevertheless, since STAT-1 is an important mediator for inflammatory signaling, rs7953280 could influence inflammation during OA progression by modulating STAT-1 binding.

### *SOCS2* deletion increases proinflammatory gene expression in response to FN-f

To assess the functional role of *SOCS2*, we used CRIS-PR-Cas9 to knock out *SOCS2* in primary human chondrocytes isolated from three individual donors. After targeting the *SOCS2* gene with two guide RNAs that flank exon 2 (a constitutive exon that contains the translational start site), we used our previously developed method that employs PCR to screen single-cell-derived colonies^53^. The screening primers generated a 1068 bp product if the region was intact and a novel 240 bp amplicon if the two guides successfully deleted the intended 828 bp region (**Fig 4A**). We saw efficient deletion in each of the three donors, with 31% of the colonies showing no deletion, 49% of the colonies showing heterozygous deletion, and 20% showing homozygous deletion (**Fig. 4B**). Sanger sequencing was used to confirm deletions, while qPCR and western blotting confirmed partial (heterozygous) or complete (homozygous) loss of *SOCS2* expression (**Supplementary Fig. 1**).

**Figure 4.**
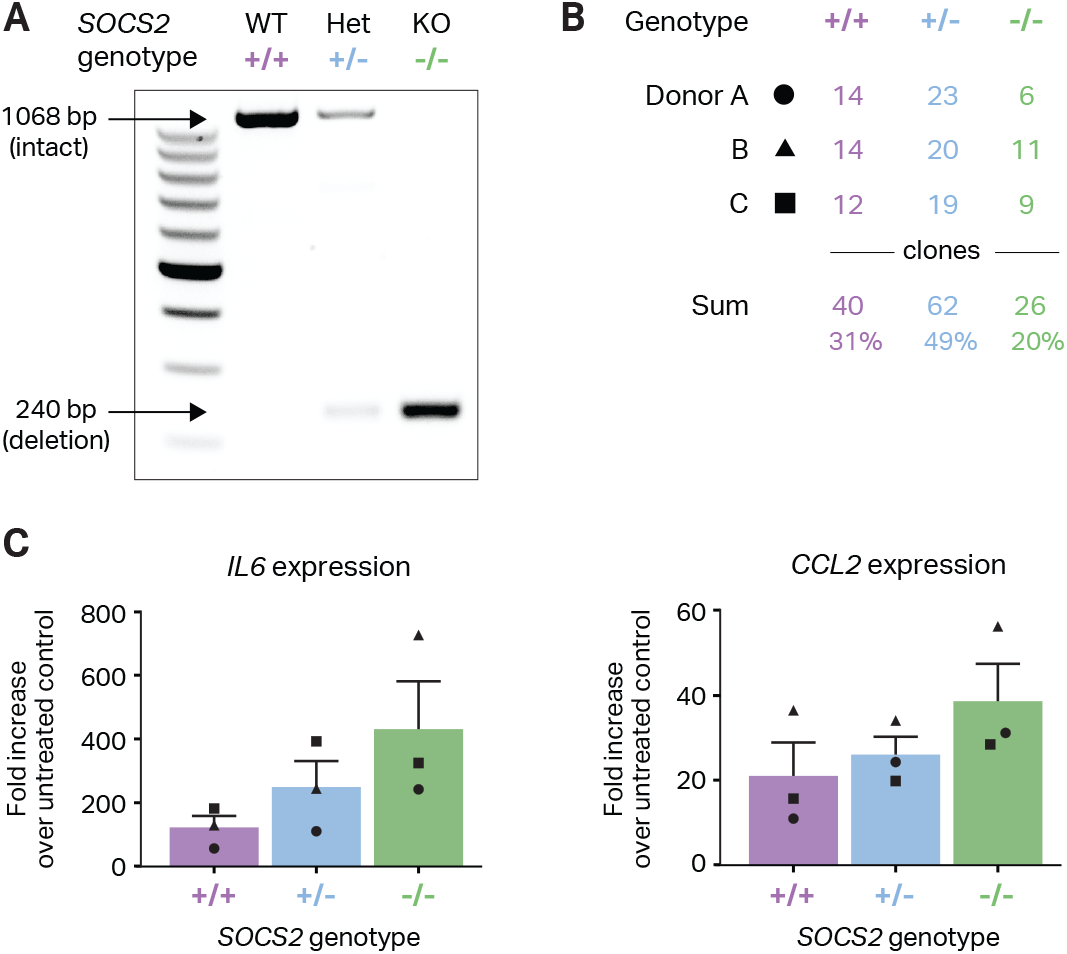
*SOCS2* deletion increases proinflammatory gene expression in response to FN-f. **(A)** PCR primers surrounding the intended SOCS2 deletion were used to screen single-cell-derived colonies from three independent donors. WT - wildtype (+/+. purple); het - heterozygous deletion (+/-, blue); KO - homozygous knockout deletion (-/-, green). **(B)** Efficiency of deleting the intended SOCS2 deletion in primary human chondrocytes from three independent donors. **(C)** qPCR at 18 hours after FN-f treatment revealed increased expression of the proinflammatory genes IL6 and CCL2 in SOCS2 deletion colonies from three independent donors.

Because *SOCS2* is a known negative regulator of the inflammatory response in other settings^48,54^, we hypothesized that *SOCS2* deletion would lead to an increased expression of inflammatory cytokines in chondrocytes during the response to FN-f. To test this hypothesis, we treated chondrocytes with defined genotypes (wild type, heterozygous, or homozygous knockout) with either FN-f or PBS for 18 hours and quantified the change in pro-inflammatory cytokines C-C Motif Chemokine Ligand 2 (*CCL2*) and Interleukin 6 (*IL6*) using qPCR. *IL6* and *CCL2* have previously been shown to exhibit increased expression after 18 hours of FN-f treatment and are also implicated in OA^11,12,55,56^. Deletion of *SOCS2* led to increased expression of both *IL6* and *CCL2* in response to FN-f treatment (**Fig. 4C**), and these increases were observed in a dose-dependent fashion, with greater increases observed in the homozygous compared to heterozygous genotypes. These results suggest that the loss of *SOCS2* may promote a heightened inflammatory response to FN-f stimulation, which is consistent with a potential role in OA.

## DISCUSSION

We used a multi-omic approach to identify putative causal SNPs and genes associated with OA risk. The efficacy of this approach was supported by the identification of previously known OA risk genes including *GDF5, SLC44A2*, and *IL11*. We generated the first maps of histone H3K27ac in primary human chondrocytes and integrated this dataset with publicly available genomic datasets to reduce thousands of OA risk GWAS variants to a small list of variants and genes for further study. By generating the first Hi-C contact map of human chondrocytes, we were able to uncover 73 previously unknown connections between OA risk variants and putative target genes. Most looped variant-gene pairs (71 of 73) skipped over the nearest gene, connecting variants to genes as far as 414 Kb away. Together, DNA looping revealed 20 unique genes, 13 of which were not identified by recent fine mapping approaches^2^ and could provide new avenues for therapeutic interventions for OA.

Among the genes identified with Hi-C, four were found to be differentially expressed in our OA model. FGFR3 and *SOCS2* have previously been implicated in OA, while Tropomoyosin 1 (TPM1) and Ral Guanine Nucleotide Dissociation Stimulator Like 1 (RGL1) have not. However, TPM1, an actin-binding protein involved in the contractile system of muscle cells and the cytoskeleton of non-muscle cells, has been shown to play roles in an inflammatory response in various cell types, such as human primary coronary artery smooth muscle cells^57^ and rod bipolar and horizontal cells in the retina^58^. RGL1, which functions as a RAS effector protein that activates GTPase by stimulating nucleotide exchange, has also been shown to modulate immune response in both vascular and immune cells^59^, and, interestingly, is downregulated in human articular chondrocytes upon treatment with interleukin-1 and oncostatin-M^60^. The functions of TPM1 and RGL1 in inflammatory responses may point to potentially undiscovered roles in osteoarthritis.

One especially intriguing gene was *SOCS2*, whose promoter is looped to an OA risk SNP within a histone H3K27ac peak ~170 Kb away. *SOCS2* is known to inhibit the JAK/STAT pathway and is induced by various pro-inflammatory cytokines such as interleukin-6, growth hormone, and tumor necrosis factor-alpha^61–63^. CRISPR-mediated deletion of *SOCS2* was associated with increased expression of *IL6* and *CCL2* in our *ex vivo* model of OA, suggesting that it may also play a role in mediating inflammation in response to cartilage matrix damage. These findings make *SOCS2* a candidate for further studies and the activation of more robust *SOCS2* expression could be a goal for future therapeutic development. The regulatory role of *SOCS2* in chondrocytes is likely to be subtle, as *SOCS2* knockout mice did not show altered OA development^64^. Because this study used a global germline deletion, other members of the inflammatory cascade may have compensated for *SOCS2* loss, and it would be interesting to learn whether the inducible loss of *SOCS2* in adult chondrocytes would generate a different result.

While further work is needed to clarify the role of rs7953280 and *SOCS2* in mediating OA risk, our multi-omic analysis suggests the following potential model. In cells harboring the non-risk variant, proinflammatory cytokines such as IL-6 and matrix damage products such as FN-f may activate AP-1 via the JAK/STAT pathway. AP-1 may then bind the enhancer at the rs7953280 locus, increase enhancer activity, and upregulate transcription of *SOCS2* via a chromatin loop between the enhancer and the *SOCS2* promoter. In cells harboring the risk allele, AP-1 binding would be decreased, impeding enhancer activation and proper upregulation of *SOCS2*. As a result, JAK/STAT signaling would remain high, resulting in prolonged or heightened inflammation and further cartilage degradation. This model, while compelling, will require further experimental investigation and validation. Moreover, future experiments are required to determine to the degree to which these findings translate from our *ex vivo* model into an *in vivo* system and/or if activation of *SOCS2* could provide therapeutic avenue for OA treatment.

In total, this work identified 56 putative OA risk genes by multi-omic data integration. Moreover, we provided the first maps of histone H3K27ac in primary human chondrocytes and the first maps of 3D chromatin contacts in chondrocytes of any type. These putative risk genes and novel epigenetic datasets will provide a foundation for future studies to investigate the genetic variants responsible for OA risk and expedite our search for better prevention and treatment of OA.

## DATA AVAILABILITY

Hi-C and CUT&RUN data can be accessed through GEO accession GSE-200345.

## FUNDING

This work was supported by NIH grants (R35-GM128645 to D.H.P., R37-AR049003 to R.F.L, and R56-AG066911 to B.O.D.) and multiple NIH training grants (T32-GM067553 for E.S.D. and N.E.K. and T32-GM007092 for E.T.). The project was also supported by the National Center for Advancing Translational Sciences (NCATS) through NIH Grant UL1TR002489 and by the UNC Thurston Arthritis Research Center through a pilot and feasibility grant. E.T. was supported by the National Science Foundation Graduate Research Fellowship Program under Grant No. DGE-2040435. Any opinions, findings, and conclusions or recommendations expressed in this material are those of the author(s) and do not necessarily reflect the views of the National Science Foundation.

## ACKNOWLEDGMENTS

We thank Jesse Raab and Thomas Vierbuchen for helpful guidance on CUT & RUN protocols, Erika Deoudes for graphic design and typesetting, and Samantha Pattenden for use of the Covaris LE220 instrument which was provided by the North Carolina Biotechnology Center Institute Development Program grant 2017-IDG-1005. We would also like to thank the Gift of Hope Organ and Tissue Donor Network, the donor families, and Dr. Susan Chubinskaya for providing normal donor tissue, as well as Dr. Pranav Mishra for donor tissue procurement and Mrs. Arnavaz Hakimiyan for technical assistance.

## METHODS

### EXPERIMENTAL METHODS

#### Primary chondrocyte isolation and culture

Primary articular chondrocytes were isolated via enzymatic digestion from human talar cartilage obtained from tissue donors, without a history of arthritis, through the Gift of Hope Organ and Tissue Donor Network (Elmhurt, IL) as previously described^12,65^. For Cut and Run, two million primary articular chondrocytes from two male donors, ages 39 and 63, were plated onto four 6cm plates in DMEM/F12 media supplemented with 10% fetal bovine serum, 1% penicillin streptomycin solution, 1% amphotericin B, and 0.04% gentamicin. For genome editing, primary chondrocytes from three male donors ages 56, 59, and 64 years were cultured in 6 or 10cm dishes at a density of approximately 70,000 cells/cm^2^ in DMEM/F12 media supplemented with 10% FBS and antibiotics.

#### Fibronectin fragment (FN-f) treatment

After serum starvation, cells were treated with either purified 42 kDa endotoxin-free recombinant FN-f (final concentration 1μM in PBS), prepared as previously described, or PBS as control^15^. Cells were harvested and crosslinked after 90 min or 180 min and immediately subjected to Cut & Run, described below.

#### Hi-C

C-28/I2 cells were cultured in DMEM/F12 media with 10% fetal bovine serum, 1% penicillin streptomycin solution, 1% amphotericin B, and 0.04% gentamicin. Cells were treated with DMEM/F12 media with 1% ITS-Plus for 48 hours prior to experiments to promote the chondrocyte phenotype. Cells were then washed with 1X PBS and treated with trypsin-EDTA (0.25%) for 3 minutes. Trypsin was quenched and cells were pelleted at 4°C for 5 minutes at 300*g*. Cells were resuspended in 1mL DMEM/F12 per million cells and crosslinked in 1% formaldehyde for 10 min with rotation before quenching in a final concentration of 0.2M glycine for 5 min with rotation. Cells were pelleted at *300g* for 5 min at 4°C. Pellets were washed with cold PBS and aliquoted into ~3 million cell aliquots. Pellets were flash frozen in liquid nitrogen and stored at −80°C. In situ Hi-C was performed as previously described^66^. A full description of our methods is provided in the Supplemental Materials.

#### Hi-C data processing

In situ Hi-C datasets were processed using a modified version of the Juicer Hi-C pipeline (https://github.com/EricSDavis/dietJuicer) with default parameters as previously described ^67^. Reads were aligned to the hg19 human reference genome with bwa (v0.7.17) and MboI was used as the restriction enzyme. Four biological replicates were aligned and merged for a total of 2,779,816 Hi-C read pairs in C-28/I2 cells yielding 2,373,892,594 valid Hi-C contacts (85.40%). For visualization, the merged Hi-C contact matrix was normalized with the “KR” matrix balancing algorithm as previously described ^68^ to adjust for regional background differences in chromatin accessibility.

Looping interactions were called with Significant Interaction Peak (SIP) caller^40^ (v1.6.2) and Juicer tools (v1.14.08) using the replicate-merged, mapq > 30 filtered hic file with the following parameters: “-norm KR -g 2.0 -min 2.0 -max 2.0 -mat 2000 -d 6 -res 5000 -sat 0.01 -t 2000 -nbZero 6 -factor 1 -fdr 0.05 -del true -cpu 1 -isDroso false”. Loop anchors were expanded to 20 Kb and loops with overlapping anchors were filtered out (14 loops). This resulted in 9,271 loops after filtering.

#### Cut and Run

Primary chondrocytes were washed with 1X PBS and treated with trypsin-EDTA (0.25%) for 5 minutes. Trypsin was quenched and cells were pelleted at 4°C for 5 minutes at 1000*g*. Cells were resuspended in 1mL plain DMEM per million cells and crosslinked in 1% formaldehyde for 10 min with rotation before quenching in a final concentration of 125 mM glycine for 5 min with rotation. Cells were pelleted by spinning at 1000*g* for 5 min at 4°C. Each 2 million cell pellet was washed in 1 mL cold PBS prior to flash freezing in liquid nitrogen. We performed Cut and Run following existing protocols^69^ but modified for crosslinked cells. A full description of our methods is provided in the Supplemental Materials.

#### Cut and Run data processing and peak calling

Adaptors and low-quality reads were trimmed from paired-end reads using Trim Galore! (v0.4.3). Reads were aligned to the hg19 genome with BWA mem (v0.7.17) and sorted with Samtools (v1.9). Duplicates were removed with PicardTools (v2.10.3) and mitochondrial reads were removed with Samtools idxstats. Samtools was also used to merge donors, and index BAM files. Peaks were called from the merged alignments using MACS2 with the following settings: -f BAM -q 0.01 -g hs --nomodel --shift 100 --extsize 200 --keep-dup all -B --SPMR (v2.1.1.20160309). Peaks were then merged using bedtools (v2.26), and multicov was used to extract counts from each replicate BAM file. Signal tracks were made from alignments using deeptools (v3.0.1).

### GENOME EDITING OF CHONDROCYTES

#### Preparation of gRNA: Cas9 RNP complex

Two custom *SOCS2* Alt-R crRNAs TGACAAGGGCCTAT-TCCCAC and TTACGCATTCCCAAGGACCC were synthesized by Integrated DNA technologies (IDT). Both sequences are written 5’ to 3’ and do not include PAM sequence. The first crRNA targets the plus strand and the second the minus strand. Ribonucleoprotein (RNP) complexes containing the Cas9 enzyme and sequence-targeting guide RNAs were prepared according to the manufacturer’s recommendation. Briefly, Alt-R tracrRNA (1072533, IDT) and crRNA were resuspended in Tris-ED-TA buffer to 100 μM concentration and equimolar concentration of crRNA and tracrRNA was combined, heated at 95 °C for 5 min and cooled to room temperature to produce the gRNA. Separate RNP complex for each guide was prepared by combining the gRNA (50 μM) with Alt-R^®^ Cas9 Nuclease (61 μM) (1081058, IDT) and PBS at a ratio of 1:1.1:2 μl at room temperature for 15 min.

#### Transfection of primary human chondrocytes with RNP complex and single cell colony selection

Chondrocytes were trypsinized, washed with PBS and transfected with the RNP complex as previously described with modifications; volumes were scaled up for transfection of more cells in larger cuvettes^53^. Two million cells were resuspended in 100 μl of P3 Primary Cell Nucle-ofector™ solution (V4XP-3024, Lonza). The RNP complex and Alt-R^®^ Cas9 Electroporation Enhancer (1075916, IDT) was added to the cells. The mixture was gently pipetted up and down and transferred to 100 μl Nucleocuvette vessels (V4XP-3024, Lonza) and transfected using program ER-100 on a 4D-Nucleofector™ Core unit (Lonza). Cells were kept at room temperature for 8 minutes and then incubated in prewarmed antibiotic free media containing 20% FBS for recovery. An aliquot of the transfected cells was placed in a 96-well and used for DNA extraction and PCR. Following confirmation of editing, the transfected bulk cells were seeded at low cell density (200 cells per 6 cm^2^ dish) for generation of single-cell colonies. Individual colonies were picked under a microscope (EVOS FL, ThermoFisher), the colony was disrupted by pipetting and split into 96- and 24-well plates for genetic analysis and continued expansion, respectively.

#### PCR screening of genome-edited bulk and single-cell derived colonies

DNA was extracted using QuickExtract™ DNA Extraction Solution (Lucigen), depending on confluency 25 to 100 μl of solution was added to the wells containing the cells for 15 minutes at 37C, cell suspension was transferred to tubes and vortexed for a minute. Samples were then heated at 65 °C for 6 minutes, and 98 °C for 2 minutes. The extracted DNA solution was stored at −20 °C. PCR amplification was performed by adding 4 or 5 μl of template DNA, 1 μM forward (*SOCS2* _F1: accaagtttgtgtgggtgct) and reverse (*SOCS2*_R1: cttccagcgtgctaagaagc) primers, and EconoTaq PLUS GREEN 2X Master Mix (Lucigen) in a 25 μl reaction. PCR conditions included an initial denaturation at 94 °C for 2 minutes, 35 cycles of denaturation at 94 °C for 30 seconds, annealing at 63 °C for 30 seconds, and extension at 72 °C for 65 seconds, followed by a final extension at 72 °C for 10 minutes. Following amplification of column purified genomic DNA, the PCR product was cleaned up and sequenced using the primers described above and the Bioedit software was used to visualize the chromatograms.

#### Fibronectin fragment (FN-f) treatment and qPCR analysis of genome edited samples

Single cell colonies in 24-well plates were passaged to 6-well plates for expansion. Following genotype confirmation, colonies with similar genotype were combined and seeded at 250,000 cells per well in a 12-well plate. Cultured cells were made serum free and treated with FN-f 1 μM or PBS. Following treatment, media was removed and cells were immediately lysed in the RLT buffer. RNA was isolated with RNeasy Plus columns (Qiagen) and reverse transcribed to cDNA using qScript™ XLT cDNA SuperMix (VWR) or iScript cDNA Synthesis Kit (1708891, Bio-Rad). DNase treatment was used for the second and third donors in order to confirm that detectable *SOCS2* signal in knockout cells was due to the presence of genomic DNA. To evaluate the effect of *SOCS2* editing on inflammatory gene response quantitative polymerase chain reaction (qPCR) was performed on a QuantStudio™ 6 Flex machine (Applied Biosystems) with TaqMan™ Universal Master Mix and TaqMan Gene Expression Assays for human *CCL2* (Hs00234140_m1), *IL6* (Hs00174131_ m1), and housekeeping gene *TBP* (Hs00427620_m1). *SOCS2* expression was assessed in pooled colonies with TaqMan Gene Expression Assay Hs00919620_m1.

#### Western Blot analysis

Following genotype identification by PCR, cells from a wildtype, heterozygous and knockout colony were expanded in chondrocyte media supplemented with 5 ng/ ml bFGF and 1 ng/ml TGF-β1 (Life technologies) for 11 days. Cells were lysed in standard cell lysis buffer (1X) (Cell signaling technology) containing phenylmethanesulfonyl fluoride (PMSF; Sigma-Aldrich)and phosphatase inhibitor mix. Protein (15 μg) was separated by SDS-PAGE and transferred to nitrocellulose membrane. After blocking in 5% nonfat milk in TBST, the blot was incubated with *SOCS2* antibody (PA5-17219; 1:1000; Thermo Fisher) overnight at 4C and secondary antibody solution for 1 hour. The membrane was incubated in Radiance Plus Chemiluminescent Substrate (Azure Biosystems) and signal detected using the Azure c600 gel imaging system. The membrane was striped and incubated with the loading control beta tubulin antibody.

### DATASETS

#### Osteoarthritis GWAS

Genome-wide association statistics for 11 osteoarthritis phenotypes and lead variants identified in Boer et al.^2^ were obtained from the Musculoskeletal Knowledge Portal^70^.

#### Epigenome Roadmap Data

Consolidated reference human epigenomes for 98 cell/ tissue types were obtained from the NIH Roadmap Epigenomics Project^71^ and The Encyclopedia of DNA Elements (ENCODE) project^72^. Processed narrowPeak files for H3K27ac, H3K4me1, and H3K4me3 and BigWig files for H3K27ac were used for each cell/tissue type. Additional narrowPeak files for H3K9ac, H3K9me3, H3K27me3, and H3K36me3 were obtained for mesenchymal stem cell derived chondrocyte cultured cells (E049).

#### RNA-seq time course of fibronectin fragment (FN-f) treatment

RNA-seq data from a prior study of FN-f treated human chondrocytes was obtained from KSM Reed et al.^12^ and vst-normalized, centered, and replicate-combined. The 0-hour FN-f treatment time point was created by combining the 9 PBS-treated replicates. Genes were considered differential with a BH-adjusted p-value of 0.01 and a log2 fold-change threshold > 1.25 across any time point.

### COMPUTATIONAL METHODS

#### Cell type enrichment for OA risk variants

To identify the cell types that likely mediate genetic OA risk, we performed SNP enrichment analysis using GREGOR (Genomic Regulatory Elements and Gwas Overlap algoRithm)^19^. Publicly available H3K27ac, H3K4me1, and H3K4me3 ChIP-seq narrowPeaks files from the NIH Roadmap Epigenomics Mapping Consortium were merged and sorted using bedtools (v2.29.2)^73^ to define regulatory loci for 98 cell types in hg19. GREGOR was used to determine each cell type’s enrichment for 104 OA lead SNPs^2^ by comparing the observed overlap between regulatory loci and SNPs with their expected overlap and evaluating significance. Expected overlap is determined using a matched control set of ~500 variants that control for the number of LD proxies, gene proximity and minor allele frequency. Reference data from 1000 Genomes Phase 1 version 2 EUR panel were used with GREGOR to control for LD proxies (1Mb, r^2^ > 0.7)^74^. Results were imported into R (v4.1.0)^75^ and visualized with ggplot2^76^ and plotgardener^77^.

#### Putative OA risk variants

LD proxies for 104 OA GWAS signals from Boer et al. were identified using the 1000 Genomes European reference panel since the GWAS data primarily analyzed individuals of European ancestry (11 of 13 cohorts are of European descent). r^2^ values were calculated with the -ld function in PLINK 1.9^74,78^ using a window of 1 Mb for LD calculation. Putative OA risk variants were defined as those in high LD (r^2^> 0.8, n = 1,259) with lead variants.

#### Multi-omic integration for assigning SNPs to putative OA risk genes

We took a multi-omic approach to identify putative SNP-gene pairs implicated in OA. SNPs that 1) were predicted to affect coding regions of genes, 2) overlapped gene promoters, or 3) overlapped a regulatory peak looped to a gene’s promoter were assigned to the “Coding gene”, “Gene promoter”, or “Loops to gene promoter” categories, respectively. Genes in each category that change in response to FN-f (p ≤ 0.01 and LFC at any time point ≥ 1.25) were highlighted as putative OA risk genes.

Coding SNP-gene pairs were identified using ENSEMBL’s Variant Effect Predictor (VEP) tool. Putative OA risk variants (n = 1,259) were annotated with their predicted consequence on coding sequence using VEP run with the GRCh37.p13 human genome and default parameters. SNPs with a predicted consequence of “missense” or “synonymous” were paired with their affected genes assigned to the “Coding gene” category.

Promoter regions were defined as 2000 bp upstream and 200 bp downstream of the TSS of transcripts obtained with the TxDb.Hsapiens.UCSC.hg19.knownGene Bioconductor package for a total of 8,2960 transcripts. Gene symbols were linked to transcript ranges using the OrganismDbi and Homo.sapiens packages. Transcripts without gene symbols or those not present in the FN-f RNA-seq data were filtered out, leaving a total of 62,590 transcript promoters.

Chondrocyte regulatory regions were defined by combining Roadmap Epigenomics data with data from primary human articular chondrocytes. Specifically, H3K4me1, H3K4me3, H3K9ac, H3K9me3, H3K27ac, H3K27me3, and H3K36me3 peaks from mesenchymal stem cell derived chondrocyte cultured cells (E049) obtained through AnnotationHub (v3.1.7, snapshot date 2021-10-20)^79^ were combined with Donor-merged H3K27ac peaks from primary human articular chondrocytes. OA SNPs were overlapped with chondrocyte regulatory regions resulting in 507 SNPs.

SNPs overlapping chondrocyte regulatory regions that also overlapped a promoter region were assigned to their affected gene and added to the “Gene promoter” category. SNPs overlapping chondrocyte regulatory regions were intersected with loop calls from Hi-C in the C-28/I2 chondrocyte cell line (see Methods on Hi-C processing and loop calling). The linkOverlaps function from the InteractionSet package was used to identify chondrocyte regulatory SNPs that are connected to promoters by loops. These SNP-gene pairs were assigned to the “Loops to gene promoter” category.

#### Motif Analysis

Tomtom (v5.4.1; release date: Sat Aug 21 19:23:23 2021 −0700) from the MEME suite was used to identify motif matches for sequences surrounding the rs7953280 variant^80^. All 7-mers surrounding rs7953280 (“GGCTTTG”, “GCTTTGA”, “CTTTGAG”, “TTTGAGG”, “TTGAGGC”, “TGAGGCA”, “GAGGCAT”) and the entire 13 bp sequence (“GGCTTTGAGGCAT”) were used to identify motif matches. Sequences were input into the online motif comparison tool and queried against the JASPAR2022_ CORE_vertebrates_non-redundant_v2 and HOCOMO-COv11_core_HUMAN_mono_meme_format motif databases. Pearson correlation coefficient was used as the motif column comparison function and the significance threshold was set to an E-value < 10; no q-value threshold was set and reverse complementing of motifs was permitted. The following command summarizes the parameters used: “tomtom -no-ssc -oc. -verbosity 1 -min-overlap 5 -mi 1 -dist pearson -evalue -thresh 10.0 -time 300 query_motifs motif_databases.”

#### Transcription factor (TF) motif binding propensity

We used SNP Effect Matrix scores (SEMs) to predict the TF binding propensity between risk and non-risk SNPs in OA. Pre-calculated SEMs for 211 TF motifs were obtained from SEMpl (https://github.com/Boyle-Lab/SEMpl) and used for scoring risk and non-risk SNP sequences^52^. Binding propensity scores were determined by generating frame-shifted K-mers covering each TF motif position for both risk and non-risk sequences. K-mers were scored against 211 TF SEMs using position-weight matrix (PWM) scoring functions from the Biostrings Bioconductor package^81^. The best scoring K-mer frame for each TF motif was used to select the binding score for risk and non-risk sequences. Scores were normalized by applying inverse-log transformation, subtracting the scrambled baseline provided with each SEM, and dividing the result by the absolute value of that baseline. TFs with positive scores are predicted to be bound while negative scores are predicted to be unbound.

## SUPPLEMENTAL FIGURES

**Figure S1.**
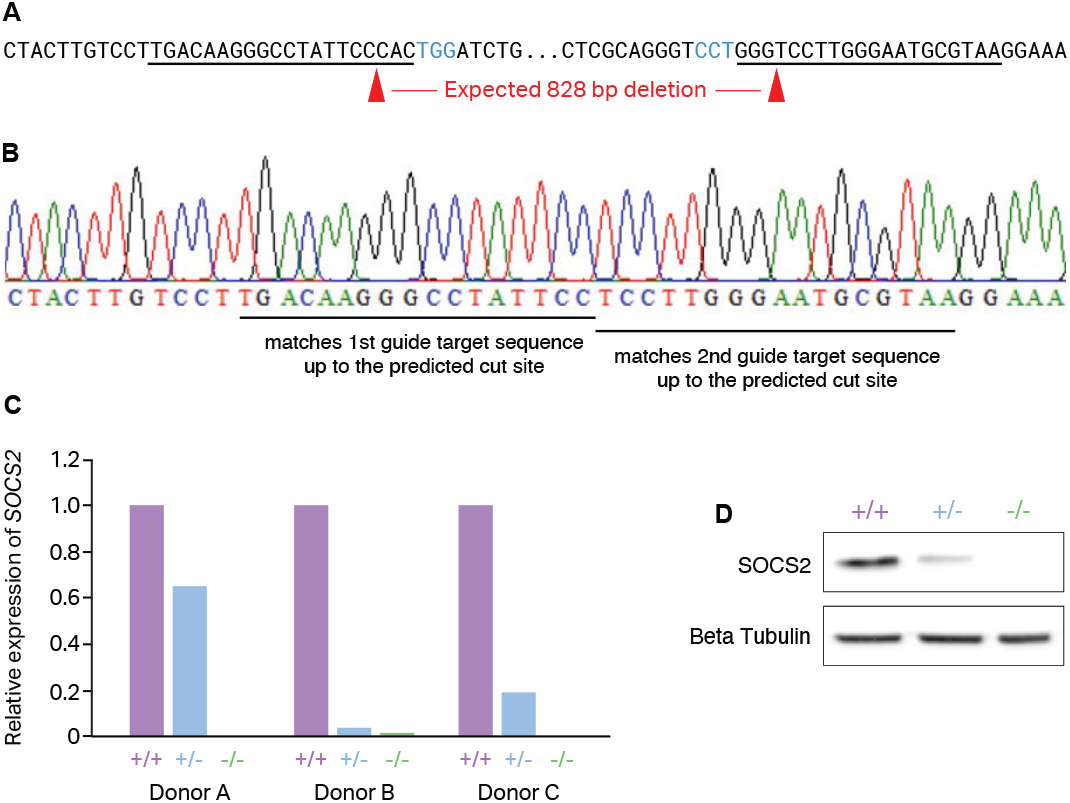
Validation of SOCS2 knockout. **(A)** Region of SOCS2 targeted by editing, PAM sites in blue, guide RNA target sites underlined, expected cut sites depicted by red triangles. **(B)** Sanger sequencing of knockout colony confirmed deletion. **(C)** SOCS2 expression was analyzed in pooled colonies by qPCR, RNA from colonies of donors A and C were treated with DNase before qPCR to eliminate any background signal of SOCS2 from remaining genomic DNA. Expression was normalized to housekeeping gene TBP and relative expression calculated. **(D)** Western blot analysis confirmed decreased expression in heterozygous colony and loss of SOCS2 in knockout colony.

## EXTENDED METHODS

### Hi-C

In situ Hi-C was performed as previously described^64^. Pellets were lysed in ice-cold Hi-C lysis buffer (10mM Tris-HCl pH 8.0, 10mM NaCl, 0.2% IGEPAL CA630) with 50μL of protease inhibitors for 15 min on ice. Cells were pelleted and washed using the same buffer. Pellets were resuspended in 50μL 0.5% SDS and incubated at 62°C for 7 min. Reactions were quenched with 145μL water and 25μL 10% Triton X-100 at 37°C for 15 min. Chromatin was digested overnight with 25μL 10X NEBuffer2 and 100U MboI at 37°C with rotation.

Reactions were incubated at 62°C for 20 min then cooled to RT. Fragment overhangs were repaired by adding 37.5μL 0.4mM biotin-14-dATP; 1.5μL each 10mM dCTP, dGTP, dTTP; 8μL 5U/μL DNA Polymerase I, Large (Klenow) Fragment and incubating at 37°C for 1.5 h with rotation. Ligation was performed by adding 673μL water, 120μL 10X NEB T4 DNA ligase buffer, 100μL 10% Triton X-100, 6μL 20mg/mL BSA, and 1μL 2000U/μL T4 DNA ligase and incubating at RT for 4 h with slow rotation. Samples were pelleted at 2500*g*, resuspended in 432μL water, 18μL 20mg/mL proteinase K, 50μL 10% SDS, and 46μL 5M NaCl, incubated at 55°C for 30 min, and then transferred to 68°C overnight.

Samples were cooled to RT and 1.6x volumes of pure ethanol and 0.1x volumes of 3M sodium acetate pH 5.2 were added to each sample, prior to incubation at −80°C for over 4-6 h. Samples were spun at max speed at 2°C for 15 min and washed twice with 70% ethanol. Pellets were dissolved in 130μL 10mM Tris-HCl pH 8.0 and incubated at 37°C for 1-2 h. Samples were stored at 4°C overnight.

DNA was sheared using the Covaris LE220 (Covaris, Woburn, MA) to a fragment size of 300-500bp in a Covaris microTUBE. DNA was transferred to a fresh tube and the Covaris microTUBE was rinsed with 70μL of water and added to the sample. A 1:5 dilution of DNA was run on a 2% agarose gel to verify successful shearing.

Sheared DNA was size selected using AMPure XP beads. 0.55x volumes of 2X concentrated AMPure XP beads were added to each reaction and incubated at RT for 5 min. Beads were reclaimed on a magnet and the supernatant was transferred to a fresh tube. 30μL 2X concentrated AMPure XP beads were added and incubated for 5 min at RT. Beads were reclaimed on a magnet and washed with fresh 70% ethanol. Beads were dried for 5 min at RT prior to DNA elution in 300μL 10mM Tris-HCl pH 8. Undiluted DNA was run on a 2% agarose gel to verify successful size selection between 300-500 bp.

150μL 10mg/mL Dynabeads MyOne Streptavidin T1 beads were washed with 400μL 1X Tween washing buffer (TWB; 250μL Tris-HCl pH 7.5, 50μL 0.5M EDTA, 10mL 5M NaCl, 25μL Tween 20, 39.675μL water). Beads were then resuspended in 300μL 2X Binding Buffer (500μL Tris-HCl pH 7.5, 100μL 0.5M EDTA, 20mL 5M NaCl, 29.4mL water), added to the DNA sample, and incubated at RT for 15 min with rotation. DNA-bound beads were then washed twice with 600μL 1X TWB at 55°C for 2 min with shaking. Beads were resuspended in 100μL 1X NEBuffer T4 DNA ligase buffer, transferred to a new tube, and reclaimed.

Sheared ends were repaired by resuspending the beads in 88μL 1X NEB T4 DNA Ligase Buffer with 1mM ATP, 2μL 25mM dNTP mix, 5μL 10U/uL NEB T4 PNK, 4uL 3U/uL NEB T4 DNA polymerase I, and 1uL 5U/uL NEB DNA polymerase 1, large (Klenow) fragment and incubating at RT for 30 min. Beads were washed two more times with 1X TWB for 2 min at 55°C with shaking. Beads were washed once with 100uL 1X NEBuffer 2, transferred to a new tube, and resuspended in 90uL 1X NEBuffer 2, 5uL 10mM dATP, and 5uL NEB Klenow exo minus, and incubated at 37°C for 30 min. Beads were washed two more times with 1X TWB for 2 min at 55°C with shaking. Beads were washed in 100uL 1X Quick Ligation Reaction Buffer, transferred to a new tube, reclaimed, and resuspended in 50uL 1X NEB Quick Ligation Reaction Buffer. 2uL NEB DNA Quick Ligase and 3uL of an appropriate Illumina indexed adapter (TruSeq nano) were added to each sample before incubating at RT for 15 minutes. Beads were reclaimed and washed twice with 1X TWB for 2 min at 55°C. Beads were washed in 100uL 10mM Tris-HCl pH 8, transferred to a new tube, reclaimed, and resuspended in 50uL 10mM Tris-HCl pH 8.

Hi-C libraries were amplified directly off T1 beads with 8 cycles in 5uL PCR primer cocktail, 20uL Enhanced PCR mix, and 25uL of DNA on beads. The PCR settings were as follows: 3 min at 95°C followed by 4-12 cycles of 20s 98°C, 15s at 60°C, and 30s at 72°C. Samples were held at 72°C for 5 min before holding at 4°C. Amplified samples were transferred to a new tube and brought to 250uL in 10mM Tris-HCl pH 8.

Beads were reclaimed and the supernatant containing the amplified library was transferred to a new tube. Beads were resuspended in 25uL 10mM Tris-HCl pH 8 and stored at −20°C. 0.7x volumes of warmed AMPure XP beads were added to the supernatant sample and incubated at RT for 5 min. Beads were reclaimed and washed with 70% ethanol without mixing. Ethanol was aspirated. Beads were resuspended in 100uL 10mM Tris-HCl pH 8, 70uL of fresh AMPure XP beads were added, and the solution was incubated for 5 min at RT. Beads were reclaimed and washed twice with 70% ethanol without mixing. Beads were left to dry and DNA was eluted in 25uL 10mM Tris-HCl pH 8. The resulting libraries were then quantified by Qubit and Tapestation. A low depth sequence was performed first using the Miniseq sequencer system (Illumina) and analyzed using the Juicer pipeline to assess quality. The resulting libraries underwent paired-end 2×150bp sequencing on an Illumina NovaSeq sequencer. Each replicate was sequenced to an approximate depth of 750 million reads. The full sequencing depth was 2.8 billion reads.

### Cut and Run

Following flash freezing, thawed pellets were resuspended in 1mL ice-cold nuclei isolation buffer (NE1 buffer; 20mM HEPES pH 7.5, 10mM KCl, 1mM MgCl2, 1mM DTT, 0.1% Triton X-100, 1X CPI added fresh) and incubated for 10 min at 4°C with rotation. Nuclear pellet was collected by centrifugation at 1000g for 5 min at 4°C, then resuspended in 1mL of ice-cold wash buffer (WB; 20mM HEPES pH 7.5, 0.2% Tween-20, 150mM NaCl, 150mM BSA, 0.5mM Spermidine, 10mM Na-Butyrate, 1X CPI added fresh). 10uL concanavalin A lectin beads washed and resuspended in binding buffer (BB; 20mM HEPES pH 7.5, 10mM KCl, 1mM CaCl2, 1mM MnCl2) were added to each sample and incubated for 10 min at RT. Beads were reclaimed and resuspended in 50uL antibody buffer (AbB; WB supplemented with 0.1% Triton X-100 and 2mM EDTA). 0.01ug/uL H3K27ac antibody in AbB was added to each sample and samples were incubated overnight at 4°C with mixing at 1000xRPM.

Beads were reclaimed and washed with 1mL triton wash buffer (TwB; WB supplemented with 0.1% Triton X-100) without mixing. Beads were reclaimed and resuspended in 50uL AbB. 2.5uL CUTANA pAG-MNase (Epicypher, #15-1016) was added and samples were incubated for 1 h at 4°C on a metal block. Beads were reclaimed and washed twice with 1mL TwB before resuspension in 100uL TwB. To digest chromatin, 2uL 100mM CaCl2 was added and samples were incubated for 30 min at 4°C on a metal block. Digestion was halted by the addition of 100uL 2X STOP buffer (340mM NaCl, 20mM EDTA, 4mM EGTA, 0.1% Triton X-100, 50ug/mL RNAse A). Samples were incubated for 20 min at 37°C to release pA-MNase cleaved fragments from nuclei). Beads were placed on a magnet and the supernatant containing DNA fragments was transferred to a new tube. To reverse crosslinks, 2uL 10% SDS and 2uL 20mg/mL proteinase K were added to each sample and incubated for 1 h at 65°C. DNA was purified using the Zymo DNA Clean & Concentrator Kit according to manufacturer’s protocols using 5 volumes of DNA binding buffer. DNA was eluted in 55uL water.

Sequencing libraries were prepared from CUT&RUN fragments using KAPA HyperPrep with library amplification kit (no. KK8504) following the manufacturer’s instructions. Post-ligation bead cleanup was performed with two rounds of 1.2X volumes of beads and DNA was eluted in a final volume of 25uL 10mM Tris-HCl pH 8. Library amplification was performed with 20uL of the adapter ligated DNA with 12 PCR cycles. One round post amplification cleanup was performed with 1.2X volumes of beads. The resulting libraries were then quantified by Qubit and Tapestation. A low depth sequence was performed first using the Miniseq sequencer system (Illumina) and analyzed using the Juicer pipeline to assess quality control. The resulting libraries underwent paired-end 2×150bp sequencing on an Illumina NextSeq sequencer.

## Notes

https://www.ncbi.nlm.nih.gov/geo/query/acc.cgi?acc=GSE200345

### Competing Interest Statement

The authors have declared no competing interest.

## REFERENCES

1. Hunter, D. J. & Bierma-Zeinstra, S. Osteoarthritis. Lancet 393, 1745–1759 (2019).

2. Boer, C. G. et al. Deciphering osteoarthritis genetics across 826,690 individuals from 9 populations. Cell 184, 6003–6005 (2021).

3. Reynard, L. N. & Barter, M. J. Osteoarthritis year in review 2019: genetics, genomics and epigenetics. Osteoarthritis Cartilage 28, 275–284 (2020).

4. Umans, B. D., Battle, A. & Gilad, Y. Where Are the Disease-Associated eQTLs? Trends Genet. 37, 109–124 (2021).

5. Claussnitzer, M. et al. FTO Obesity Variant Circuitry and Adipocyte Browning in Humans. N. Engl. J. Med. 373, 895–907 (2015).

6. Won, H. et al. Chromosome conformation elucidates regulatory relationships in developing human brain. Nature 538, 523–527 (2016).

7. Duan, A. et al. Chromatin architecture reveals cell type-specific target genes for kidney disease risk variants. BMC Biol. 19, 38 (2021).

8. Laarman, M. D. et al. Chromatin Conformation Links Putative Enhancers in Intracranial Aneurysm-Associated Regions to Potential Candidate Genes. J. Am. Heart Assoc. 8, e011201 (2019).

9. Chesi, A. et al. Genome-scale Capture C promoter interactions implicate effector genes at GWAS loci for bone mineral density. Nat. Commun. 10, 1260 (2019).

10. Loeser, R. F. Integrins and chondrocyte-matrix interactions in articular cartilage. Matrix Biol. 39, 11–16 (2014).

11. van den Bosch, M. H. J., van Lent, P. L. E. M. & van der Kraan, P. M. Identifying effector molecules, cells, and cytokines of innate immunity in OA. Osteoarthritis Cartilage 28, 532–543 (2020).

12. Reed, K. S. M. et al. Transcriptional response of human articular chondrocytes treated with fibronectin fragments: an in vitro model of the osteoarthritis phenotype. Osteoarthritis Cartilage 29, 235–247 (2021).

13. Forsyth, C. B., Pulai, J. & Loeser, R. F. Fibronectin fragments and blocking antibodies to alpha2beta1 and alpha5beta1 integrins stimulate mitogen-activated protein kinase signaling and increase collagenase 3 (matrix metalloproteinase 13) production by human articular chondrocytes. Arthritis Rheum. 46, 2368–2376 (2002).

14. Pulai, J. I. et al. NF-κB Mediates the Stimulation of Cytokine and Chemokine Expression by Human Articular Chondrocytes in Response to Fibronectin Fragments. The Journal of Immunology 174, 5781–5788 (2005).

15. Wood, S. T. et al. Cysteine-mediated redox regulation of cell signaling in chondrocytes stimulated with fibronectin fragments. Arthritis rheumatol. 68, 117–126 (2016).

16. Xie, D. L., Meyers, R. & Homandberg, G. A. Fibronectin fragments in osteoarthritic synovial fluid. J. Rheumatol. 19, 1448–1452 (1992).

17. Homandberg, G. A., Wen, C. & Hui, F. Cartilage damaging activities of fibronectin fragments derived from cartilage and synovial fluid. Osteoarthritis Cartilage 6, 231–244 (1998).

18. Homandberg, G. A. Potential regulation of cartilage metabolism in osteoarthritis by fibronectin fragments. Front. Biosci. 4, D713–30 (1999).

19. Schmidt, E. M. et al. GREGOR: evaluating global enrichment of trait-associated variants in epigenomic features using a systematic, data-driven approach. Bioinformatics 31, 2601–2606 (2015).

20. Loeser, R. F., Goldring, S. R., Scanzello, C. R. & Goldring, M. B. Osteoarthritis: a disease of the joint as an organ. Arthritis Rheum. 64, 1697–1707 (2012).

21. Sandell, L. J. & Aigner, T. Articular cartilage and changes in Arthritis: Cell biology of osteoarthritis. Arthritis Res. Ther. 3, 107 (2001).

22. Caron, M. M. J. et al. BAPX-1/NKX-3.2 Acts as a Chondrocyte Hypertrophy Molecular Switch in Osteoarthritis. Arthritis & Rheumatology vol. 67 2944–2956 (2015).

23. Pelletier, J. P., Martel-Pelletier, J. & Abramson, S. B. Osteoarthritis, an inflammatory disease: potential implication for the selection of new therapeutic targets. Arthritis Rheum. 44, 1237–1247 (2001).

24. Venetianer, P. Are synonymous codons indeed synonymous? Biomol. Concepts 3, 21–28 (2012).

25. Zeng, Z. & Bromberg, Y. Predicting Functional Effects of Synonymous Variants: A Systematic Review and Perspectives. Frontiers in Genetics vol. 10 (2019).

26. Tuerlings, M. et al. RNA Sequencing Reveals Interacting Key Determinants of Osteoarthritis Acting in Subchondral Bone and Articular Cartilage: Identification of *IL11* and *CHADL* as Attractive Treatment Targets. Arthritis & Rheumatology vol. 73 789–799 (2021).

27. Song, J. et al. MicroRNA-488 regulates zinc transporter SLC39A8/ZIP8 during pathogenesis of osteoarthritis. J. Biomed. Sci. 20, 31 (2013).

28. Jain, S., Gautam, V. & Naseem, S. Acute-phase proteins: As diagnostic tool. J. Pharm. Bioallied Sci. 3, 118–127 (2011).

29. Boeuf, S. et al. Subtractive gene expression profiling of articular cartilage and mesenchymal stem cells: serpins as cartilage-relevant differentiation markers. Osteoarthritis Cartilage 16, 48–60 (2008).

30. Wanner, J. et al. Proteomic profiling and functional characterization of early and late shoulder osteoarthritis. Arthritis Res. Ther. 15, R180 (2013).

31. Francis-West, P. H., Parish, J., Lee, K. & Archer, C. W. BMP/GDF-signalling interactions during synovial joint development. Cell Tissue Res. 296, 111–119 (1999).

32. Miyamoto, Y. et al. A functional polymorphism in the 5’ UTR of GDF5 is associated with susceptibility to osteoarthritis. Nat. Genet. 39, 529–533 (2007).

33. Southam, L. et al. An SNP in the 5’-UTR of GDF5 is associated with osteoarthritis susceptibility in Europeans and with in vivo differences in allelic expression in articular cartilage. Hum. Mol. Genet. 16, 2226–2232 (2007).

34. Capellini, T. D. et al. Ancient selection for derived alleles at a GDF5 enhancer influencing human growth and osteoarthritis risk. Nat. Genet. 49, 1202–1210 (2017).

35. Richard, D. et al. Evolutionary Selection and Constraint on Human Knee Chondrocyte Regulation Impacts Osteoarthritis Risk. Cell 181, 362–381.e28 (2020).

36. Muthuirulan, P. et al. Joint disease-specificity at the regulatory base-pair level. Nat. Commun. 12, 4161 (2021).

37. Steinberg, J. et al. A molecular quantitative trait locus map for osteoarthritis. Nat. Commun. 12, 1309 (2021).

38. Steinberg, J. et al. Decoding the genomic basis of osteoarthritis. bioRxiv 835850 (2020) doi:10.1101/835850.

39. Maurano, M. T. et al. Systematic localization of common disease-associated variation in regulatory DNA. Science 337, 1190–1195 (2012).

40. Rowley, M. J. et al. Analysis of Hi-C data using SIP effectively identifies loops in organisms from C. elegans to mammals. Genome Res. 30, 447–458 (2020).

41. Zhou, S. et al. Conditional Deletion of Fgfr3 in Chondrocytes leads to Osteoarthritis-like Defects in Temporomandibular Joint of Adult Mice. Scientific Reports vol. 6 (2016).

42. Tang, J. et al. Fibroblast Growth Factor Receptor 3 Inhibits Osteoarthritis Progression in the Knee Joints of Adult Mice. Arthritis Rheumatol 68, 2432–2443 (2016).

43. Okura, T. et al. Activated FGFR3 prevents subchondral bone sclerosis during the development of osteoarthritis in transgenic mice with achondroplasia. J. Orthop. Res. 36, 300–308 (2018).

44. Lin, A. C. et al. Modulating hedgehog signaling can attenuate the severity of osteoarthritis. Nature Medicine vol. 15 1421–1425 (2009).

45. Li, X. et al. Species-specific biological effects of FGF-2 in articular cartilage: implication for distinct roles within the FGF receptor family. J. Cell. Biochem. 113, 2532–2542 (2012).

46. Shu, C. C. et al. Ablation of Perlecan Domain 1 Heparan Sulfate Reduces Progressive Cartilage Degradation, Synovitis, and Osteophyte Size in a Preclinical Model of Posttraumatic Osteoarthritis. Arthritis & Rheumatology vol. 68 868–879 (2016).

47. de Andrés, M. C. et al. Suppressors of cytokine signalling (SOCS) are reduced in osteoarthritis. Biochem. Biophys. Res. Commun. 407, 54–59 (2011).

48. Paul, I. et al. The ubiquitin ligase Cullin5 *SOCS2* regulates NDR1/STK38 stability and NF-κB transactivation. Sci. Rep. 7, 42800 (2017).

49. Motomura, H. et al. A selective c-Fos/AP-1 inhibitor prevents cartilage destruction and subsequent osteophyte formation. Biochem. Biophys. Res. Commun. 497, 756–761 (2018).

50. Fisch, K. M. et al. Identification of transcription factors responsible for dysregulated networks in human osteoarthritis cartilage by global gene expression analysis. Osteoarthritis Cartilage 26, 1531–1538 (2018).

51. Gao, X., Sun, Y. & Li, X. Identification of key gene modules and transcription factors for human osteoarthritis by weighted gene co-expression network analysis. Exp. Ther. Med. 18, 2479–2490 (2019).

52. Nishizaki, S. S. et al. Predicting the effects of SNPs on transcription factor binding affinity. Bioinformatics 36, 364–372 (2020).

53. D’Costa, S., Rich, M. J. & Diekman, B. O. Engineered Cartilage from Human Chondrocytes with Homozygous Knockout of Cell Cycle Inhibitor p21. Tissue Eng. Part A 26, 441–449 (2020).

54. Monti-Rocha, R. et al. *SOCS2* Is Critical for the Balancing of Immune Response and Oxidate Stress Protecting Against Acetaminophen-Induced Acute Liver Injury. Front. Immunol. 9, 3134 (2018).

55. Wang, T. & He, C. Pro-inflammatory cytokines: The link between obesity and osteoarthritis. Cytokine Growth Factor Rev. 44, 38–50 (2018).

56. Wojdasiewicz, P., Poniatowski, Ł. A. & Szukiewicz, D. The role of inflammatory and anti-inflammatory cytokines in the pathogenesis of osteoarthritis. Mediators Inflamm. 2014, 561459 (2014).

57. Li, R., Liang, Y. & Lin, B. Accumulation of systematic TPM1 mediates inflammation and neuronal remodeling by phosphorylating PKA and regulating the FABP5/NF-κB signaling pathway in the retina of aged mice. Aging Cell 21, e13566 (2022).

58. Gagat, M. et al. CRISPR-Based Activation of Endogenous Expression of TPM1 Inhibits Inflammatory Response of Primary Human Coronary Artery Endothelial and Smooth Muscle Cells Induced by Recombinant Human Tumor Necrosis Factor α. Frontiers in Cell and Developmental Biology vol. 9 (2021).

59. Kirkby, N. S. et al. COX-2 protects against atherosclerosis independently of local vascular prostacyclin: identification of COX-2 associated pathways implicate Rgl1 and lymphocyte networks. PLoS One 9, e98165 (2014).

60. Yang, Y. et al. Novel role of circRSU1 in the progression of osteoarthritis by adjusting oxidative stress. Theranostics 11, 1877–1900 (2021).

61. Starr, R. et al. A family of cytokine-inducible inhibitors of signalling. Nature 387, 917–921 (1997).

62. Metcalf, D. et al. Gigantism in mice lacking suppressor of cytokine signalling-2. Nature 405, 1069–1073 (2000).

63. Santangelo, C., Scipioni, A., Marselli, L., Marchetti, P. & Dotta, F. Suppressor of cytokine signaling gene expression in human pancreatic islets: modulation by cytokines. Eur. J. Endocrinol. 152, 485–489 (2005).

64. Samvelyan, H. J., Huesa, C., Cui, L., Farquharson, C. & Staines, K. A. The role of accelerated growth plate fusion in the absence of *SOCS2* on osteoarthritis vulnerability. Bone Joint Res. 11, 162–170 (2022).

65. Loeser, R. F., Pacione, C. A. & Chubinskaya, S. The combination of insulin-like growth factor 1 and osteogenic protein 1 promotes increased survival of and matrix synthesis by normal and osteoarthritic human articular chondrocytes. Arthritis Rheum. 48, 2188–2196 (2003).

66. Rao, S. S. P. et al. A 3D map of the human genome at kilobase resolution reveals principles of chromatin looping. Cell 159, 1665–1680 (2014).

67. Durand, N. C. et al. Juicer provides a one-click system for analyzing loop-resolution Hi-C experiments. Cell Syst. 3, 95–98 (2016).

68. Knight, P. A. & Ruiz, D. A fast algorithm for matrix balancing. IMA J. Numer. Anal. 33, 1029–1047 (2013).

69. Skene, P. J. & Henikoff, S. An efficient targeted nuclease strategy for high-resolution mapping of DNA binding sites. Elife 6, (2017).

70. Kiel, D. P. et al. The Musculoskeletal Knowledge Portal: Making Omics Data Useful to the Broader Scientific Community. J. Bone Miner. Res. 35, 1626–1633 (2020).

71. Bernstein, B. E. et al. The NIH Roadmap Epigenomics Mapping Consortium. Nat. Biotechnol. 28, 1045–1048 (2010).

72. ENCODE Project Consortium. An integrated encyclopedia of DNA elements in the human genome. Nature 489, 57–74 (2012).

73. Quinlan, A. R. & Hall, I. M. BEDTools: a flexible suite of utilities for comparing genomic features. Bioinformatics 26, 841–842 (2010).

74. 1000 Genomes Project Consortium et al. A map of human genome variation from population-scale sequencing. Nature 467, 1061–1073 (2010).

75. R Core Team. R: A Language and Environment for Statistical Computing. (2021).

76. Wickham, H. ggplot2: Elegant Graphics for Data Analysis. (2016).

77. Kramer, N. E. et al. Plotgardener: Cultivating precise multi-panel figures in R. Bioinformatics (2022) doi:10.1093/bioinformatics/btac057.

78. Purcell, S. et al. PLINK: a tool set for whole-genome association and population-based linkage analyses. Am. J. Hum. Genet. 81, 559–575 (2007).

79. Morgan, M., Carlson, M., Tenenbaum, D. & Arora, S. AnnotationHub: Client to access AnnotationHub resources. R package version 2, (2017).

80. Gupta, S., Stamatoyannopoulos, J. A., Bailey, T. L. & Noble, W. S. Quantifying similarity between motifs. Genome Biol. 8, R24 (2007).

81. Pagès, H., Aboyoun, P., Gentleman, R. & DebRoy, S. Biostrings: Efficient manipulation of biological strings. (2021).

